# SARS-like coronaviruses in horseshoe bats (*Rhinolophus* spp.) in Russia, 2020

**DOI:** 10.1101/2021.05.17.444362

**Authors:** Sergey V. Alkhovsky, Sergey V. Lenshin, Alexey V. Romashin, Tatyana V. Vishnevskaya, Oleg I. Vyshemirsky, Yulia I. Bulycheva, Dmitry K. Lvov, Asya K. Gitelman

## Abstract

We found and genetically described two novel SARS-like coronaviruses in feces and oral swabs of the great (*R. ferrumequinum*) and the lesser (*R. hipposideros*) horseshoe bats in southern region of Russia. The viruses, named Khosta-1 and Khosta-2, together with related viruses from Bulgaria and Kenya, form a separate phylogenetic lineage. We found an evidence of recombination events in evolutionary history of Khosta-1, which involved the acquisition of structural proteins S, E, and M as well as nonstructural genes ORF3, ORF6, ORF7a, and ORF7b from a virus that is closely related to Kenyan isolate BtKY72. Examination of bats by RT-PCR revealed that 62,5% of great horseshoe bats in one of the caves were positive for Khosta-1 virus while its overall prevalence was 14%. The prevalence of Khosta-2 was 1,75%. Our results show that SARS-like coronaviruses circulate in horseshoe bats in the region and provide a new data on their genetic diversity.

## Introduction

Horseshoe bats (*Rhinolophidae: Rhinolophus*) are considered as a main natural reservoir and source of zoonotic coronaviruses (CoV) which caused epidemic outbreak of severe acute respiratory syndrome (SARS) and COVID-19 pandemic in 2002 and 2019, respectively (1,2). These viruses, designated as SARS-CoV and SARS-CoV-2, together with related viruses found in bats and other animals (SARS-like coronaviruses or SARS-CoV-like viruses) belong to the subgenus *Sarbecovirus* of the genus *Betacoronavirus* of the family *Coronaviridae* (3). Horseshoe bats are wide distributed in Asia, Europe, and North Africa. In East Asia (in particular, in People’s Republic of China (PRC)) SARS-CoV-like viruses circulate in multiple rhinolophid’s species, however the Chinese rufous (*R. sinicus*), the greater (*R. ferrumequinum*), the intermediate (*R. affinis*), and the king (*R. rex*) horseshoe bats seem to be of major importance (4). In Europe SARS-CoV-like viruses was found in the greater, the lesser (*R. hipposideros*), the Miditerranean (*R. euryale*), Mehely’s (*R. mehelyi*), and Blasius’s (*R. blasii*) horseshoe bats (5–8). The prevalence of SARS-like coronaviruses among bats in different caves-colonies can vary from 0% to 60% (4,7,9,10). In Russia three species of horseshoe bats (the greater, the lesser, and Miditerranean) are common in southern regions, lying below about 44° north latitude, mostly including North Caucasus and Crimea. In present work we hypothesized that SARS-like coronaviruses circulate in the region in local populations of horseshoe bats. To test this hypothesis, we examined the colonies of bats located in the southern macroslope of the Greater Caucasus on northern coast of the Black Sea in Russia. Using metagenomic analysis we found and genetically described two new SARS-like coronaviruses in feces and oral swabs of the greater and the lesser horseshoe bats. Further PCR analysis showed a high degree of infection of bats with discovered viruses in some locations.

## Materials and methods

The samples from bats were collected in Sochi National Park (Sochi-Adler, Krasnodar krai, Russia) and surrounding areas in March-October 2020. The Sochi National Park is located on southern macroslope of the Greater Caucasus descending to the northern coast of the Black Sea (Figure 1). The park stuffs keep records of more than 300 karst formations (caves, breaks, mines, clefts, etc.) that are natural refuges for bats and other troglophilous animal species. Bats were caught by hand in eight locations including five caves as well as basements and attics of houses (Table 1). The bats were caught in the frame of ongoing surveillance of bats populations constantly carried out in the park. The species of the animals was determined based on their morphological characters by an experienced park zoologist. The length of the forearm and weight of animals were measured. To collect bat oral swabs (saliva and buccal cells) an urological swab was placed in the bat’s mouth for 10-15 sec and then placed in 250 μl of phosphate-buffered saline (PBS). To collect feces, an animal was placed in a small white cotton bag for 10-15 min. After that the animal was released, and the feces were collected from the walls of the bag into cryovials. No bats were harmed during sample collection. A total of 120 samples of oral swabs and 77 samples of feces from five species of bats were collected (Table 1). The samples were delivered to the laboratory on ice and stored at −70°C until start of research.

**Figure 1.**
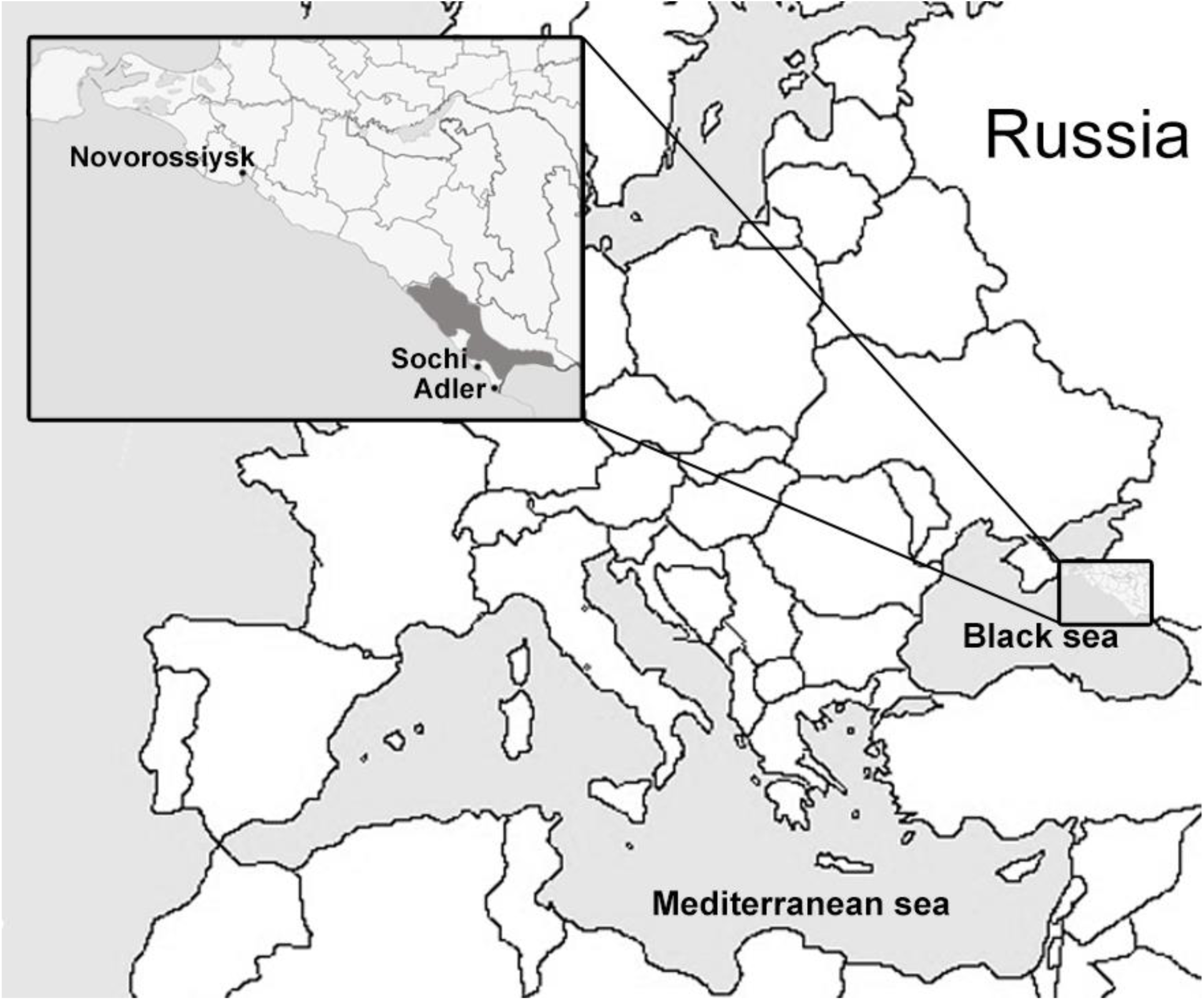
Map of the region where bat samples were collected. Location of Sochi National Park and surrounding area is shown in grey.

**Table 1.**
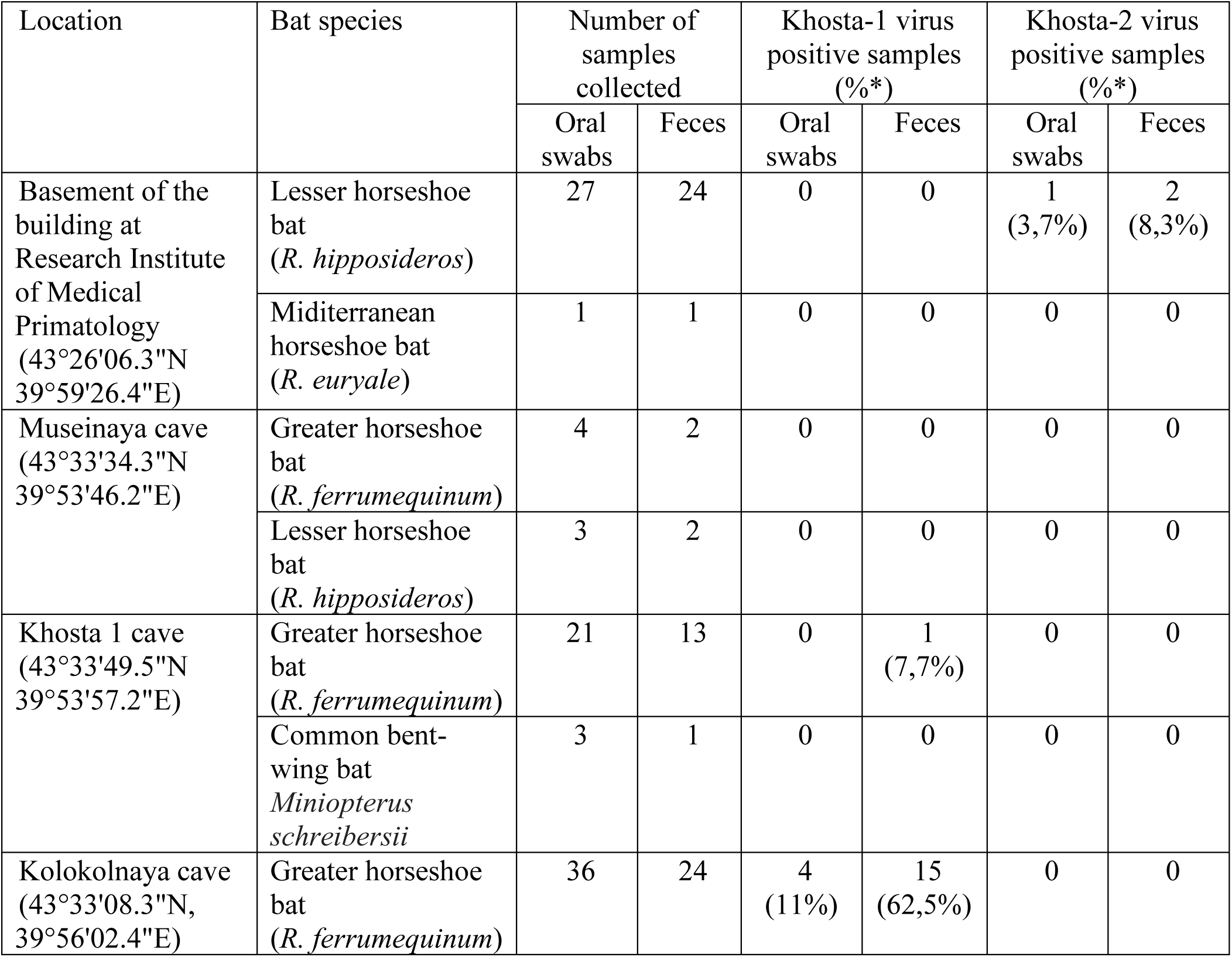
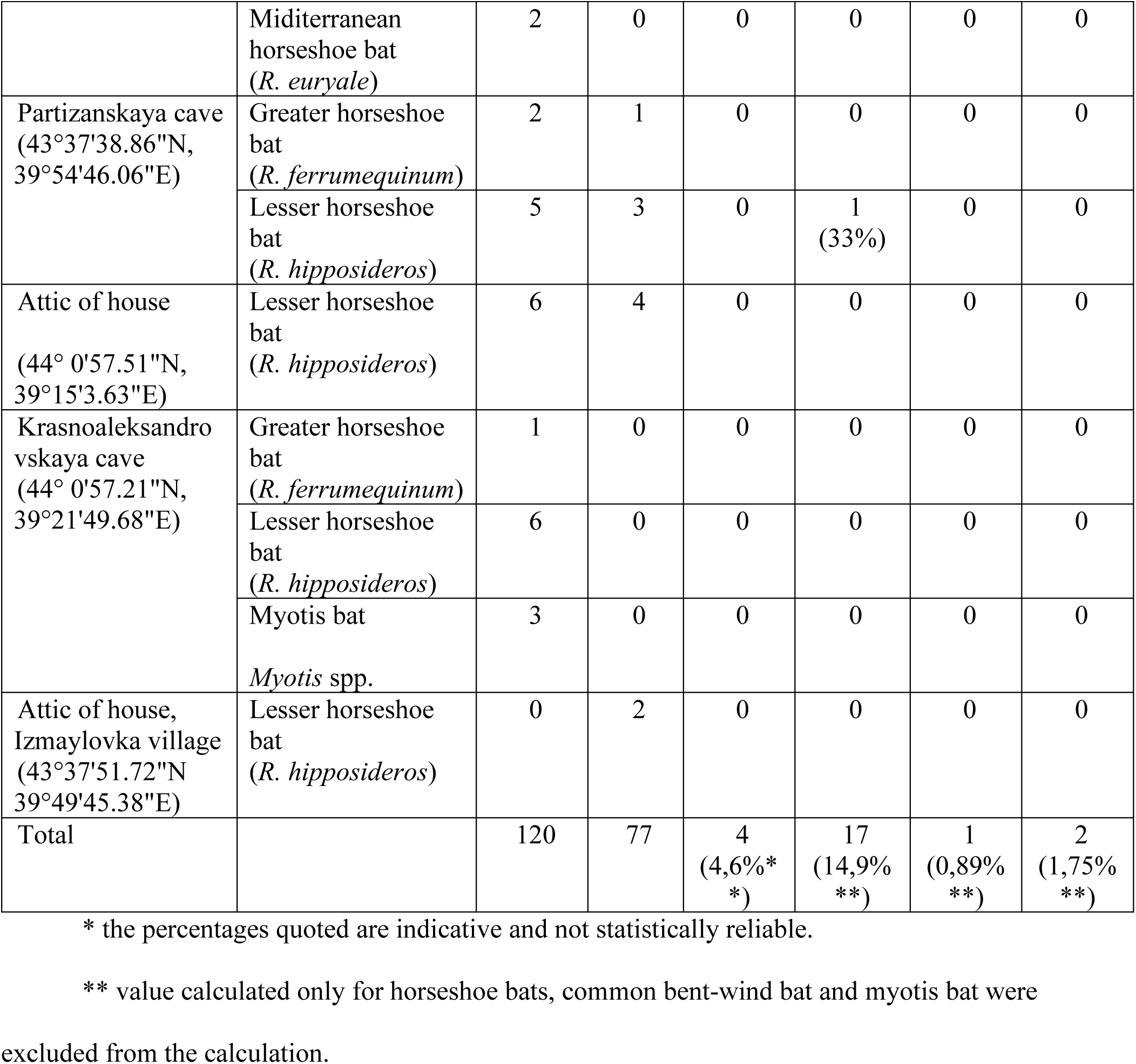
The bat samples collected and results of RT-PCR testing for Kosta-1 and Khosta-2 viruses.

Metagenomic analysis was conduct as described anywhere. Briefly, the feces were suspended and homogenized in 0,5 ml of PBS and pooled by 0,1 ml in three samples (feces from 20-30 animals in the pool). Pooled samples were clarified by centrifugation (10 000 g, 15 min) and treated with DNase I and RNase I_f_ (NEB, Great Britain) for removing of naked out-capsid nucleic acid. The viral particles were sedimented from treated samples by ultracentrifugation (30 000 g, 1 h) through 2 ml of 20% sucrose. Virus plaque was resuspended in 0,5 ml PBS. Total RNA was isolated from 0,25 ml of obtained solution with TRIzol LS reagent (ThermoFisher Scientific, USA).

Total RNA from oral swabs were isolated with TRIzol LS reagent from individual samples and pooled by 20 μl in five pooled samples (20-25 samples in the pool). The pooled RNA was precipitated by isopropyl alcohol with addition of glycogen followed by additional clarification with RNeasy MinElute Cleanup kit (Qiagen, Germany). NEBNext rRNA Depletion kit (NEB) was used to remove bacterial and eukaryotic rRNA from total RNA isolated from pooled samples. Treated RNA was used for cDNA library preparation by NEBNext Ultra II RNA library kit for Illumina (NEB).

The libraries were sequenced on a HiSeq4000 instrument (Illumina, USA) at the facility of Resource Center “BioBank” of the Research Park of Saint Petersburg State University (Saint-Petersburg, Russia). Reads were filtered by quality, trimmed to remove adapter’s sequences, and assembled *de novo* using CLC Genomics Workbench 7.0 software (Qiagen). Obtained contig sequences were analyzed using blastx algorithm by DIAMOND software (11) against nr ‘Viruses’ database that included all reference viral sequences available in GenBank at December 2020. Nucleotide and deduced amino acid sequences were alignment by ClustalW implemented in MEGAX software (12). The best substitution model was evaluated for each alignment by the Model selection module in MEGAX software. Phylogenetic trees were inferred by ‘maximum likelihood’ method using appropriate model with 1000 bootstrap replicates by MEGAX software. Similarity plot analysis was conducted by SimPlot software (13). Possible recombination was analyzed by RDP5 software (14).

Primers and probes for specific detection of discovered coronaviruses were developed based on obtained sequences by Beacon 7.0 software (Premier Biosoft, USA). Real-time RT-PCR was conducted with TaqPath 1-Step Multiplex Master Mix (ThermoFisher Scientific, USA) and total RNA isolated by TRIzol LS reagent from individual oral swabs and feces samples.

## Results

### Results of sequencing of the samples

In total, 124 522 978 reads for three pooled fecal samples and 170 112 341 reads for five pooled oral swab samples were obtained. The reads were *de novo* assembled in contigs and analyzed by blastx algorithm for presence of viral sequences. The search results revealed two extended contigs with a length of approximately 29 Kb with open reading frames (ORFs) with similarity to members of the genus *Betacoronavirus* in the pools one and three of fecal samples, respectively. With further analysis, a near-complete genome of two novel SARS-like coronaviruses was sequenced. Matching contigs have also been found in corresponding oral swab samples, but with smaller length and coverage. Two found SARS-like coronaviruses were named BtCoV/Khosta-1/Rh/Russia/2020 and BtCoV/Khosta-2/Rh/Russia/2020 and placed in GenBank by accession numbers MZ190137 and MZ190138, respectively. Bellow they are referred to as Khosta-1 and Khosta-2, respectively.

### Genetic and phylogenetic analysis

The genomic organization of Khosta-1 and Khosta-2 is similar to that of other SARS-like coronaviruses (Figure 2). Approximately two thirds of the genome of coronaviruses is occupied by ORF1a and ORF1b genes which encode the proteins of the replicative complex and translated as ORF1ab polyprotein due to ribosomal shifting. The rest of the genome contains genes of structural proteins (S, E, M, and N), which form a virion, as well as several non-structural proteins (ORF3, ORF6, ORF7, ORF8, ORF9, and ORFX), the presence and structure of which varies in different viruses (3). Genome organization of Khosta-1 and Khosta-2 has the greatest similarity with BtCoV/BM48-31/2008 and BtKY72 viruses – two SARS-like coronaviruses found in horseshoe bats in Bulgaria and Kenya in 2008 and 2007, respectively (8,15). Their peculiarity is the absent of ORF8 gene which is common in bat SARS-like coronaviruses from East and Southeast Asia (Figure 2).

**Figure 2.**
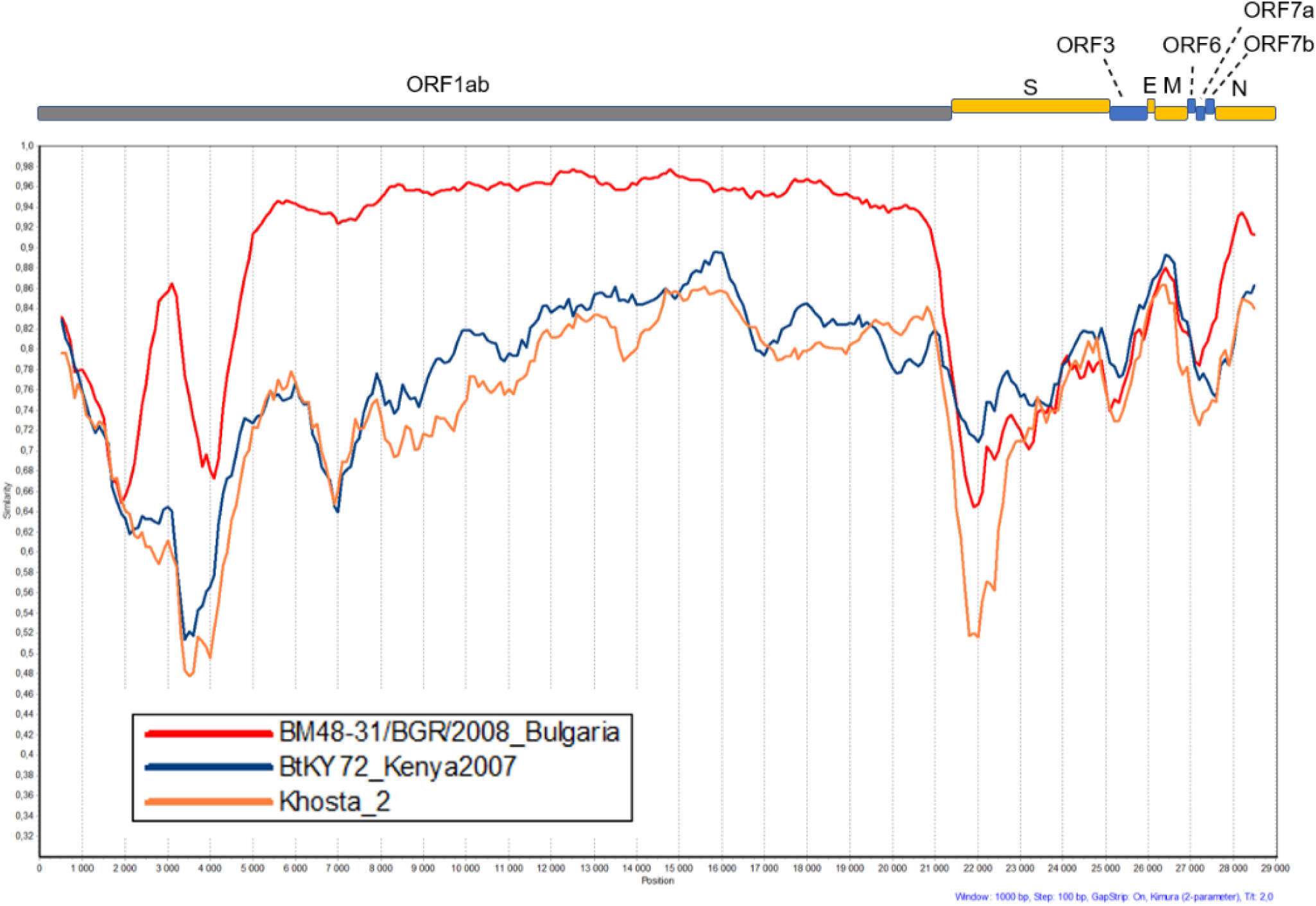
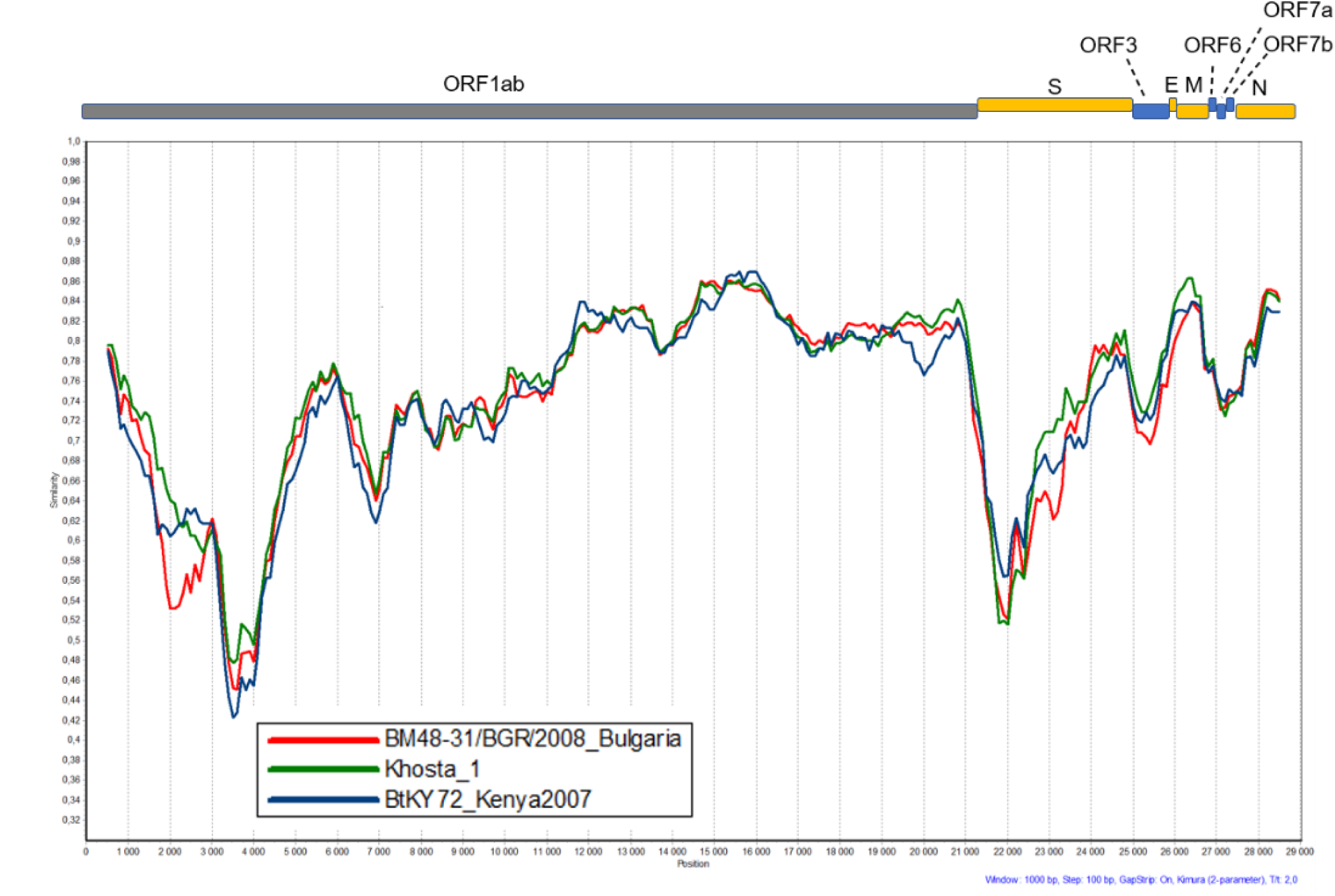
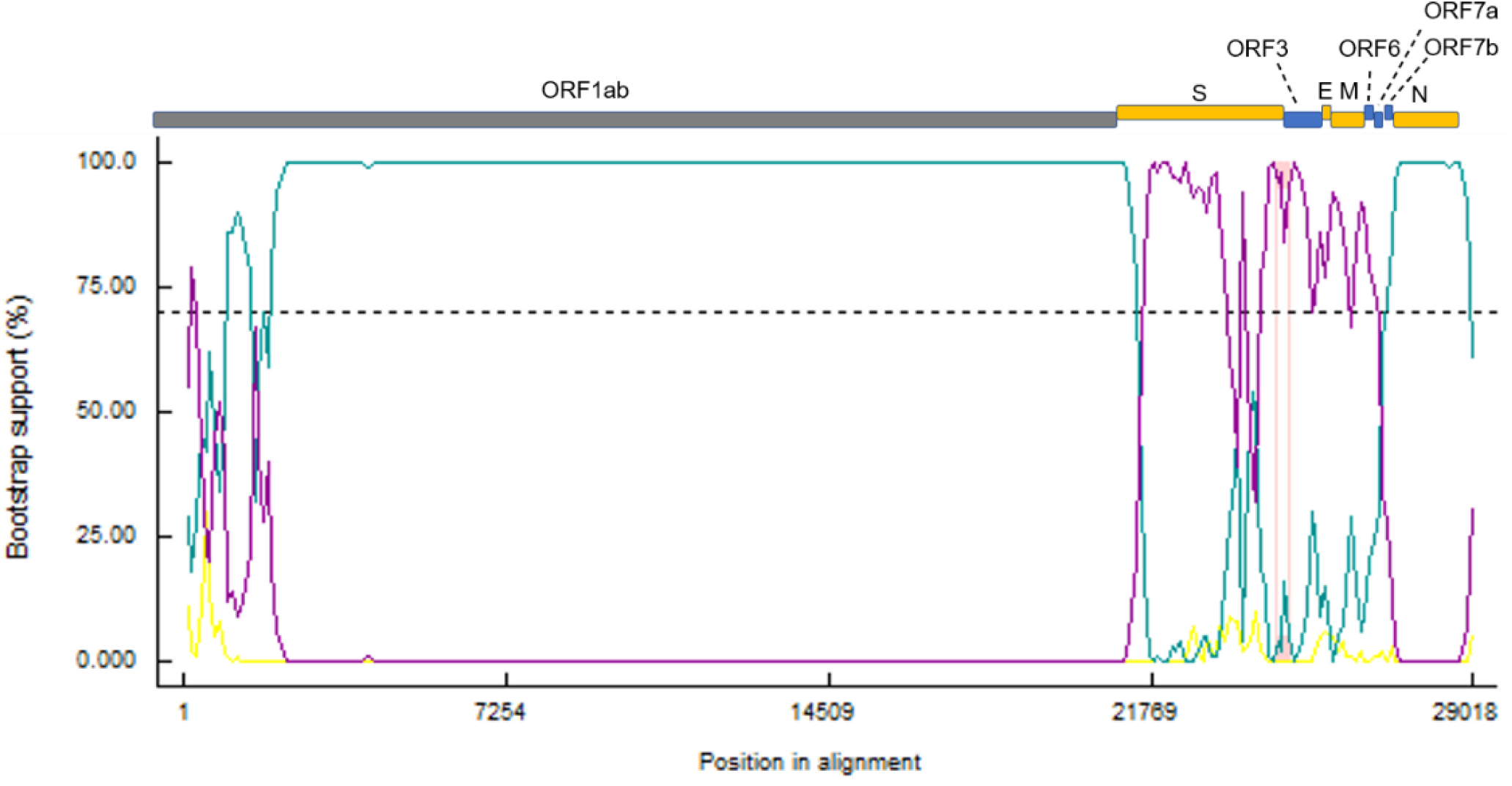
Scheme of the genome of Khosta-1 and Khosta-2 viruses with the designation of the main ORFs, and results of Simplot and recombination analysis. **A**. Khosta-1 was used as a query sequence and Khosta-2, BM48-31/BGR/2008, and BtKy72 viruses were used as reference sequences. **B**. Khosta-2 was used as a query sequence and Khosta-1, BM48-31/BGR/2008, and BtKy72 were used as reference sequences. The analysis was performing with Kimura (2-parametr) model, window size of 1000 bases and a step size of 100 bases. **C.** Results of bootstrap analysis of recombination events in Khosta-1 genome by RDP5 software.

Pairwise alignments of the deduced proteins of Khosta-1 and Khosta-2 virus with those of other SARS-like coronaviruses showed its highest similarity with BtCoV/BM48-31/2008 and BtKY72 viruses (Table 2). Khosta-1 is closest related to BtCoV/BM48-31/2008 with 92,5% aa and 99% aa identity in the conservative ORF1a and ORF1b proteins, respectively. Similarity of Khosta-1 with SARS-CoV and related viruses from China are on average 81,5% aa identity in ORF1a protein and 96% in ORF1b protein. Comparison Khosta-1 with SARS-CoV-2 viruses revealed 77,5% and 94,2% aa identity for ORF1a and ORF1b proteins, respectively. Despite the high similarity of Khosta-1 and BtCoV/BM48-31/2008 in ORF1a and ORF1b proteins, structural proteins S, E, and M of Khosta-1 are more similar to those of Kenyan virus BtKY72. Khosta-1 and BtKY72 share 89,1%, 98,7%, and 97,29% aa identity for S, E, and M proteins, whereas these values for Khosta-1 and BtCoV/BM48-31/2008 are 84,37%, 89,47, and 95%, respectively. N protein of Khosta-1 is more similar to that of BtCoV/BM48-31/2008 (96,64%aa identity) than BtKY72 (92,6% aa identity).

**Table 2.**
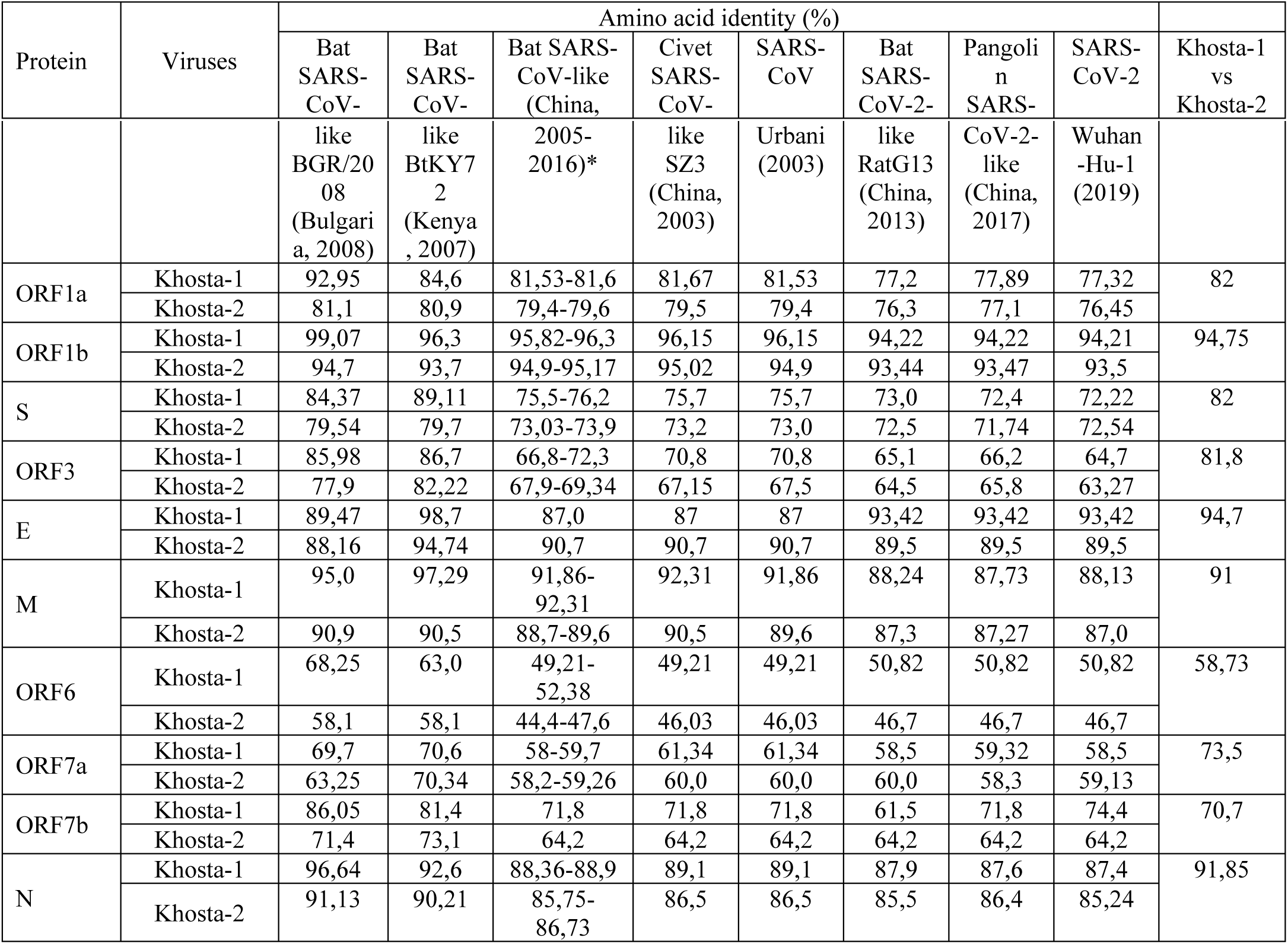
Similarity (%) of deduced amino acid sequences of proteins of Khosta-1 and Khosta-2 viruses with certain representatives of *Sarbecovirus* subgenus (lineage B of betacoronaviruses).

| Protein | Viruses | Amino acid identity (%) |  |  |  |  |  |  |  | Khosta-1<br>vs<br>Khosta-2 |
| --- | --- | --- | --- | --- | --- | --- | --- | --- | --- | --- |
|  |  | Bat SARS-CoV- | Bat SARS-CoV- | Bat SARS-CoV-like<br>(China, | Civet SARS-CoV- | SARS-CoV | Bat SARS-CoV-2- | Pangolin SARS- | SARS-CoV-2 |  |

|  |  | like<br>BGR/20<br>08<br>(Bulgari<br>a, 2008) | like<br>BtKY7<br>2<br>(Kenya<br>, 2007) | 2005-2016)* | like<br>SZ3<br>(China,<br>2003) | Urbani<br>(2003) | like<br>RatG13<br>(China,<br>2013) | CoV-2-like<br>(China,<br>2017) | Wuhan<br>-Hu-1<br>(2019) |  |
| --- | --- | --- | --- | --- | --- | --- | --- | --- | --- | --- |
| ORF1a | Khosta-1 | 92,95 | 84,6 | 81,53-81,6 | 81,67 | 81,53 | 77,2 | 77,89 | 77,32 | 82 |
|  | Khosta-2 | 81,1 | 80,9 | 79,4-79,6 | 79,5 | 79,4 | 76,3 | 77,1 | 76,45 |  |
| ORF1b | Khosta-1 | 99,07 | 96,3 | 95,82-96,3 | 96,15 | 96,15 | 94,22 | 94,22 | 94,21 | 94,75 |
|  | Khosta-2 | 94,7 | 93,7 | 94,9-95,17 | 95,02 | 94,9 | 93,44 | 93,47 | 93,5 |  |
| S | Khosta-1 | 84,37 | 89,11 | 75,5-76,2 | 75,7 | 75,7 | 73,0 | 72,4 | 72,22 | 82 |
|  | Khosta-2 | 79,54 | 79,7 | 73,03-73,9 | 73,2 | 73,0 | 72,5 | 71,74 | 72,54 |  |
| ORF3 | Khosta-1 | 85,98 | 86,7 | 66,8-72,3 | 70,8 | 70,8 | 65,1 | 66,2 | 64,7 | 81,8 |
|  | Khosta-2 | 77,9 | 82,22 | 67,9-69,34 | 67,15 | 67,5 | 64,5 | 65,8 | 63,27 |  |
| E | Khosta-1 | 89,47 | 98,7 | 87,0 | 87 | 87 | 93,42 | 93,42 | 93,42 | 94,7 |
|  | Khosta-2 | 88,16 | 94,74 | 90,7 | 90,7 | 90,7 | 89,5 | 89,5 | 89,5 |  |
| M | Khosta-1 | 95,0 | 97,29 | 91,86-92,31 | 92,31 | 91,86 | 88,24 | 87,73 | 88,13 | 91 |
|  | Khosta-2 | 90,9 | 90,5 | 88,7-89,6 | 90,5 | 89,6 | 87,3 | 87,27 | 87,0 |  |
| ORF6 | Khosta-1 | 68,25 | 63,0 | 49,21-52,38 | 49,21 | 49,21 | 50,82 | 50,82 | 50,82 | 58,73 |
|  | Khosta-2 | 58,1 | 58,1 | 44,4-47,6 | 46,03 | 46,03 | 46,7 | 46,7 | 46,7 |  |
| ORF7a | Khosta-1 | 69,7 | 70,6 | 58-59,7 | 61,34 | 61,34 | 58,5 | 59,32 | 58,5 | 73,5 |
|  | Khosta-2 | 63,25 | 70,34 | 58,2-59,26 | 60,0 | 60,0 | 60,0 | 58,3 | 59,13 |  |
| ORF7b | Khosta-1 | 86,05 | 81,4 | 71,8 | 71,8 | 71,8 | 61,5 | 71,8 | 74,4 | 70,7 |
|  | Khosta-2 | 71,4 | 73,1 | 64,2 | 64,2 | 64,2 | 64,2 | 64,2 | 64,2 |  |
| N | Khosta-1 | 96,64 | 92,6 | 88,36-88,9 | 89,1 | 89,1 | 87,9 | 87,6 | 87,4 | 91,85 |
|  | Khosta-2 | 91,13 | 90,21 | 85,75-86,73 | 86,5 | 86,5 | 85,5 | 86,4 | 85,24 |  |

In contrast, Khosta-2 does not have such an increased similarity with some group of sarbecoviruses and has 79-81% aa identity with SARS-CoV viruses and 76-77% with SARS-CoV-2 viruses in ORF1a protein. ORF1b protein of Khosta-2 has 93,5-95% aa identities with all other bat SARS-like coronaviruses. Comparison of proteins of Khosta-1 and Khosta-2 showed that these viruses differ from each other at about the same level as Khosta-2 differs from other bat SARS-like coronaviruses (Table 2).

### Recombination analysis

Genome sequence similarity between Khosta-1, Khosta-2, BtCoV/BM48-31/2008, and BtKY72 was analyzed by Simplot and RDP5 software (Figure 2A-2C). Simplot analysis showed high degree of similarity of Khosta-1 and BtCoV/BM48-31/2008 in the ORF1ab and N genes and a decrease in the similarity in the S-ORF7b region. Results of analysis carried out by RDP5 software using different methods implemented in the program showed clear signals of recombination events in evolutionary history of Khosta-1 (Figure 2C). Recombination events presumably included acquisition of S-ORF7b region by ancestor of Khosta-1 virus from a virus closely related to BtKY72.

### Phylogenetic analysis

Phylogenetic analysis based on ORF1ab protein sequences showed that Khosta-1, Khosta-2, BtCoV/BM48-31/2008, and BtKY72 form a monophyletic lineage located between SARS-CoV and SARS-CoV-2 lineages of the *Sarbecovirus* subgenus (Figure 3A). The separate cluster this group of viruses also formed on phylogenetic tree based on nucleotide sequences of S gene (Figure 3B). Topology of tree confirms the probable recombination event in evolutionary history of Khosta-1. In ORF1ab tree Khosta-1 is grouped together with BtCoV/BM48-31/2008, while in S gene tree with BtKY72.

**Figure 3.**
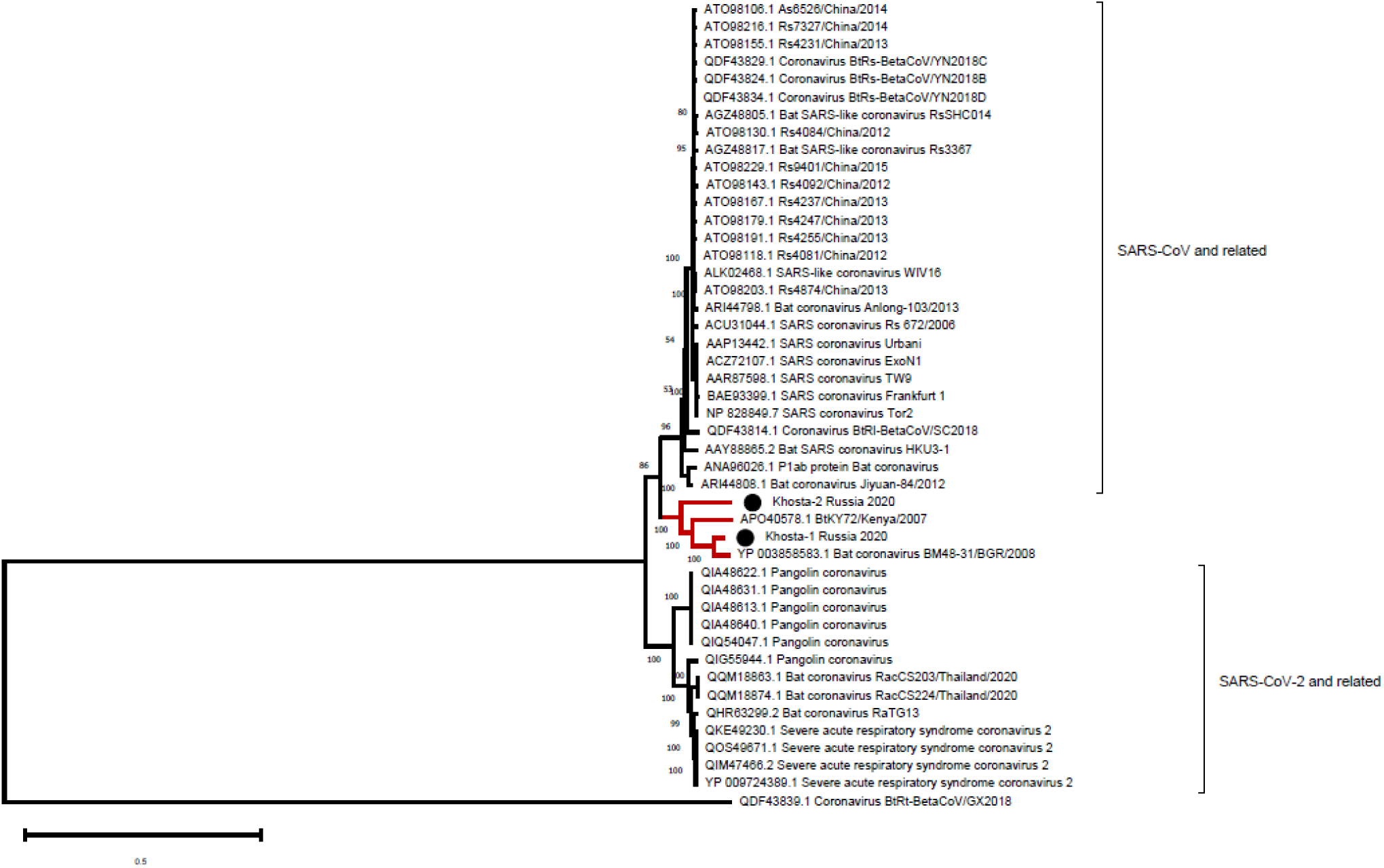
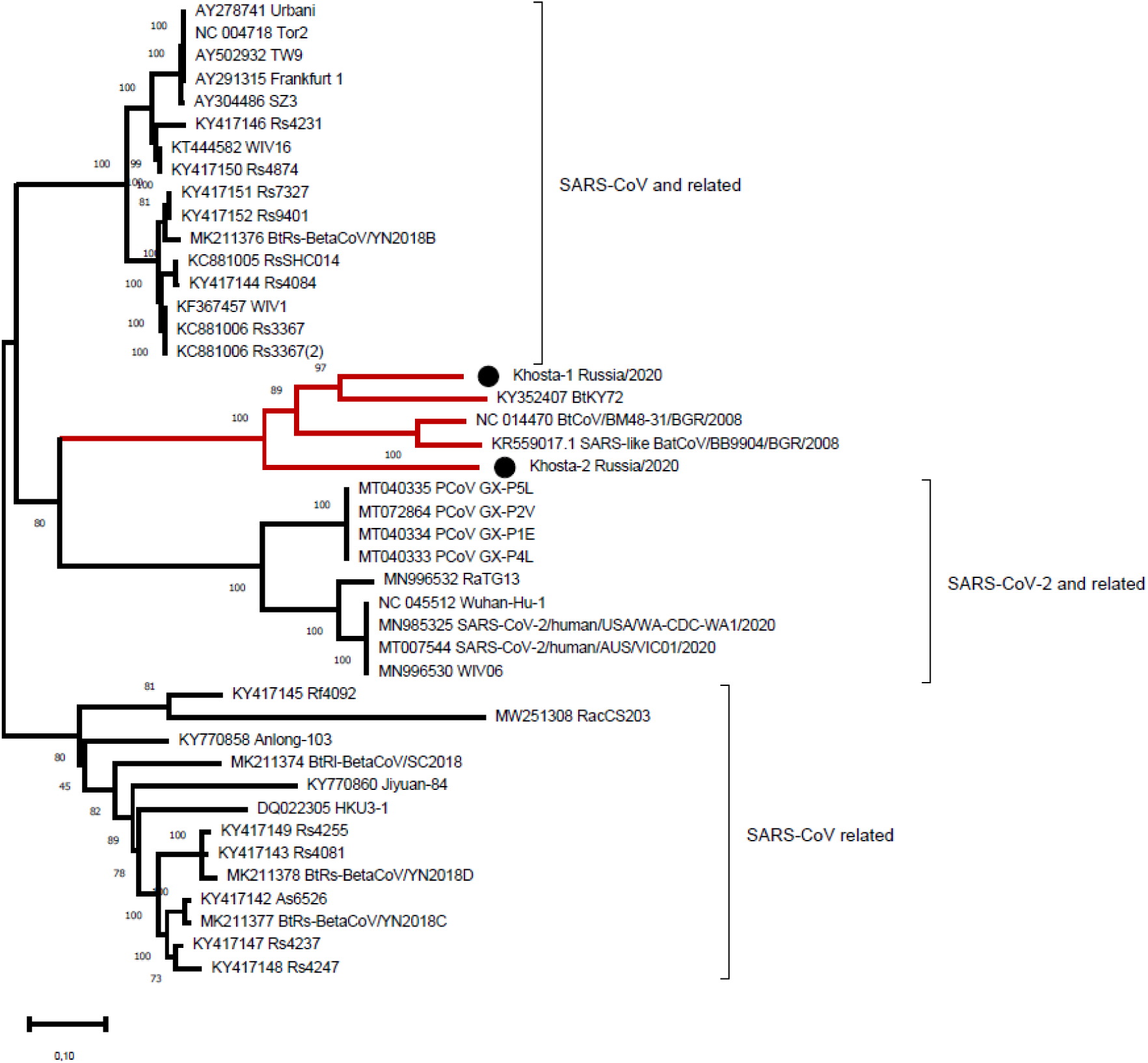
Phylogenetic trees inferred by maximum likelihood method based on analysis of amino acid sequences of ORF1ab protein (**A**) and nucleotide sequences of S gene (**B**). HKU9-like strain Bt-BetaCoV/GX2018, belonged to subgenus *Nobecovirus* (lineage D), was used as an outer group for ORF1ab protein phylogeny. The viruses described in present work are marked by black circle.

### Analysis of receptor binding motif (RBM) of S protein

Alignment of amino acid sequences of RBM of Khosta-1 and Khosta-2 with certain sarbecoviruses are presented in Figure 4. This is a highly variable region where multiple substitutions and deletions occur among SARS-CoV related viruses. Khosta-1 and Khosta-2 as well as BtCoV/BM48-31/2008 have a common deletion of four aa in the N-part of RBM. This deletion partially overlapping with the deletion that is characteristic of HKU3-1 and related strains of bat SARS-CoV-like viruses that unable to bind angiotensin-converting enzyme 2 (ACE2) receptor. We analyzed aa positions in RBM which are thought to be crucial for binding of ACE2 receptor and, therefore, important for adaptation of bat SARS-like coronaviruses to human (16,17). Only at position 442 Khosta-1 and Khosta-2 share a common amino acid (L), which is also inherent in SARS-CoV-2 and related viruses. Despite the significant genetic distance between Khosta-1 and BtKY72, crucial positions in RBM and their context coincide between them. In contrast, position 479, 480, and 487 of Khosta-2 coincides poorly with other groups of viruses (Figure 4).

**Figure 4.**
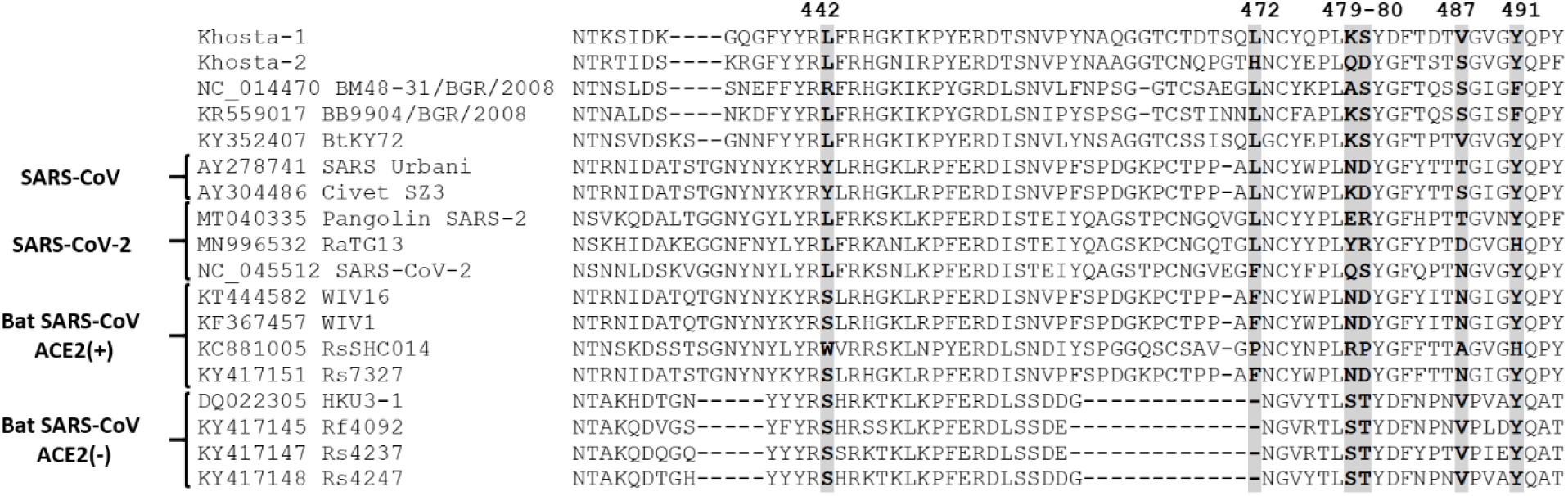
Alignment of receptor binding motif (RBM) of receptor binding domain (RBD) of S protein of Khosta-1 and Khosta-2 with certain sarbecoviruses. Position (442, 472, 479, 480, 487, 491 numbering by SARS-CoV Urbani) in RBM which are thought to be important for adaptation SARS-CoV-like viruses to human ACE2 receptor (16,17) are bolded. Bat SARS-CoV-like viruses that are capable or not capable of utilizing ACE2 receptor are marked with “ACE2(+)” or “ACE2(-)”, respectively.

### PCR testing

We developed primers and probes for specific detection of Khosta-1 and Kosta-2 viruses in feces and oral swabs by real time RT-PCR (Appendix Table). Results of PCR testing of samples are presented in Table 1. RNA of Khosta-1 was detected mostly in the greater horseshoe bats collected in Kolokolnaya cave. All four found positive oral swabs belonged to animals with positive fecal samples. In other locations Khosta-1 virus was detected only in two fecal samples – from the greater horseshoe bat from Khosta 1 cave and the lesser horseshoe bat from Partizanskaya cave, respectively. Also note that RNA of Khosta-1 was detected in feces much more often than in oral swabs and in a higher titer (based on Ct value, data not shown). RNA of Khosta-2 virus was detected in two lesser horseshoe bats collected in basement of the building at Research Institute of Medical Primatology. In one animal RNA of Khosta-2 was detected in both feces and oral swabs, and in another only swab.

## Discussion

The emergence of SARS-CoV and SARS-CoV-2 viruses is a result of adaptation of SARS-CoV-like viruses, circulated in the horseshoe bats, to human (18,19). Horseshoe bats are widespread and, presumably, SARS-like coronaviruses circulate in all parts of their range including Asia, Europe and northern Africa. However, little information exists on the genetic diversity of bat SARS-like coronaviruses in the regions outside East and Southeast Asia. We described here two novel SARS-like coronaviruses circulated in the horseshoe bats in southern region of Russia. Khosta-1 and Khosta-2 viruses are closely related to viruses recently described in Bulgaria (strain BtCoV/BM48-31/2008) and Kenya (strain BtKY72) (8,15). Together they form a separate “western” (as they are found in the western part of the range of horseshoe bats) phylogenetic lineage of bat SARS-like coronaviruses. A feature of these viruses is the absence of ORF8 gene, which is common in SARS-CoV, SARS-CoV-2, and most of bat SARS-like coronaviruses of eastern lineages.

SASR-CoV and SARS-CoV-2 recognize host angiotensin-converting enzyme 2 (ACE2) as its receptor. Crucial for binding of ACE2 receptor amino acids (442, 487, 479, 487, and 491) are located in the RBM of the S protein (17,20). These amino acids and its context in RBM of Khosta-1 and Khosta-2, like most other bat SARS-like coronaviruses, are quite different from SASR-CoV and SARS-CoV-2 viruses. The most bat SARS-like coronaviruses unable to bind ACE2 receptor of human and do not infect its cells (21). However, several strains of bat SARS-like coronaviruses that can use ACE2 receptor and whose proteins are highly similar to SARS-CoV or SARS-CoV-2 have been recently found in the Chinese rufous (*R. sinicus*) and the intermediate (*R. affinis*) horseshoe bats in People’s Republic of China (22–25). In this vein, it is argued that direct progenitors of SARS-CoV and SARS-CoV-2 originated after sequential recombination events between bat SARS-like coronaviruses. Since the recombination is of great importance in the evolution of coronaviruses, we analyzed possible recombination events in the western lineage of bat SARS-like coronaviruses. Despite the small number of known full-length sequences (only four), we observe an evidence of recombination in evolutionary history of Khosta-1. The alleged recombination event involved the acquisition of structural proteins S, E, and M as well as nonstructural genes ORF3, ORF6, ORF7a, and ORF7b from a virus that is closer to the Kenyan isolate BtKY72 than to the European strain BtCoV/BM48-31/2008. Based on this, we can assume that genetic diversity of viruses in the region is significantly higher than is known today and there is constant exchange of genes between them. These findings require further investigation of the diversity of circulating variants, with particular emphasis on the diversity of the S gene.

Using RT-PCR we showed that 14% of tested horseshoe bats were carriers for Khosta-1 virus and 1,75% for Khosta-2 virus. However, most of the Khosta-1 positive samples were found in only one cave (Kolokolnaya cave) where infection rate of greater horseshoe bats reached 62,5%. This bias, together with the small number of samples from other locations, makes it difficult to accurately estimate the prevalence of Khosta-1 in the region and requires further research. The closest European region where such studies have been carried out is Bulgaria: according to the data obtained by J. Drexler with coauthor (2010) SARS-like coronaviruses were detected in 13,3% of greater horseshoe bat, 15,9% of Blasius’s horseshoe bat, 30,8% of Mehely’s horseshoe bat, and 32,1% of Miditerranean horseshoe bat (8). Another studies found 38.8% positive lesser horseshoe bats in Slovenia and 37,9% positive greater horseshoe bats in France (5,6). All this data show that the prevalence of SARS-like coronaviruses in horseshoe bats in their western part of range can vary widely between different species, locations, and possibly the time of year of observation.

In conclusion, we have shown that SARS-like coronaviruses circulate in horseshoe bats in southern region of Russia and provide a new information on its genomics. Genetic diversity, prevalence, host range, as well as potential threat to human of these viruses remain to be determined.

## Acknowledgments

The work was funded by Russian Foundation for Basic Research (RFBR), project No. 20-04-60154.

## Author Bio

(first author only, unless there are only 2 authors)

Sergey V. Alkhovsky is a head of laboratory of biotechnology at D.I. Ivanovsky Institute of Virology of N.F. Gamleya National Center on Epidemiology and Microbiology in Moscow, Russia. His scientific interests are ecology, genomics, and evolution of zoonotic viruses with special emphasizing to emerging and reemerging viruses that pose serious threat to public health. It includes arboviruses as well as zoonotic viruses of rodents, bats, and birds, distributed in Northern Eurasia.

## References

1. Lau SK, Woo PC, Li KS, Huang Y, Tsoi HW, Wong BH, et al. Severe acute respiratory syndrome coronavirus-like virus in Chinese horseshoe bats. Proc Natl Acad Sci U S A. 2005;102(39):14040–5.

2. Li W, Shi Z, Yu M, Ren W, Smith C, Epstein JH, et al. Bats are natural reservoirs of SARS-like coronaviruses. Science. 2005; 310(5748):676–9.

3. de Groot RJ, Baker SC, Baric R, Enjuanes L, Gorbalenya AE, Holmes K V, et al. Family Coronaviridae. In: King AM, Adams MJ, Carstens EB, Lefkowitz EJ, editors. Virus taxonomy : classification and nomenclature of viruses : ninth report of the International Committee on Taxonomy of Viruses. 1sted ed. London: Elsevier; 2012. p. 806–28.

4. Fan Y, Zhao K, Shi Z-L, Zhou P. Bat Coronaviruses in China. Viruses. 2019;11(3):210.

5. Rihtarič D, Hostnik P, Steyer A, Grom J, Toplak I. Identification of SARS-like coronaviruses in horseshoe bats (Rhinolophus hipposideros) in Slovenia. Arch Virol. 2010;155(4):507–14.

6. Ar Gouilh M, Puechmaille SJ, Diancourt L, Vandenbogaert M, Serra-Cobo J, Lopez Roïg M, et al. SARS-CoV related Betacoronavirus and diverse Alphacoronavirus members found in western old-world. Virology. 2018;517:88–97.

7. Balboni A, Palladini A, Bogliani G, Battilani M. Detection of a virus related to betacoronaviruses in Italian greater horseshoe bats. Epidemiol Infect. 2011;139(2):216–9.

8. Drexler JF, Gloza-Rausch F, Glende J, Corman VM, Muth D, Goettsche M, et al. Genomic Characterization of Severe Acute Respiratory Syndrome-Related Coronavirus in European Bats and Classification of Coronaviruses Based on Partial RNA-Dependent RNA Polymerase Gene Sequences. J Virol. 2010;84(21):11336–49.

9. Wang LF, Shi Z, Zhang S, Field H, Daszak P, Eaton BT. Review of bats and SARS. Emerg Infect Dis. 2006;12(12):1834–40.

10. Drexler JF, Corman VM, Drosten C. Ecology, evolution and classification of bat coronaviruses in the aftermath of SARS. Antiviral Res. 2014;101:45–56.

11. Buchfink B, Xie C, Huson DH. Fast and sensitive protein alignment using DIAMOND. Nature Methods. 2015.12(1):59–60.

12. Tamura K, Peterson D, Peterson N, Stecher G, Nei M, Kumar S. MEGA5: molecular evolutionary genetics analysis using maximum likelihood, evolutionary distance, and maximum parsimony methods. Mol Biol Evol. 2011;28(10):2731–9.

13. Lole KS, Bollinger RC, Paranjape RS, Gadkari D, Kulkarni SS, Novak NG, et al. Full-length human immunodeficiency virus type 1 genomes from subtype C-infected seroconverters in India, with evidence of intersubtype recombination. J Virol. 1999;73(1):152–60.

14. Martin DP, Murrell B, Golden M, Khoosal A, Muhire B. RDP4: Detection and analysis of recombination patterns in virus genomes. Virus Evol. 2015;1(1):vev003.

15. Tong S, Conrardy C, Ruone S, Kuzmin I V, Guo X, Tao Y, et al. Detection of novel SARS-like and other coronaviruses in bats from Kenya. Emerg Infect Dis. 2009;15(3):482–5.

16. Li F. Receptor recognition and cross-species infections of SARS coronavirus. 2013;100(1):246–54.

17. Wan Y, Shang J, Graham R, Baric RS, Li F. Receptor Recognition by the Novel Coronavirus from Wuhan: an Analysis Based on Decade-Long Structural Studies of SARS Coronavirus. J Virol. 2020;94(7):e00127.

18. Li W, Shi Z, Yu M, Ren W, Smith C, Epstein JH, et al. Bats are natural reservoirs of SARS-like coronaviruses. Science. 2005;310(5748):676–9.

19. Zhou P, Yang X Lou, Wang XG, Hu B, Zhang L, Zhang W, et al. A pneumonia outbreak associated with a new coronavirus of probable bat origin. Nature. 2020; 579(7798):270–273.

20. Li W, Moore MJ, Vasllieva N, Sui J, Wong SK, Berne MA, et al. Angiotensin-converting enzyme 2 is a functional receptor for the SARS coronavirus. Nature. 2003 Nov 27;426(6965):450–4.

21. Ren W, Qu X, Li W, Han Z, Yu M, Zhou P, et al. Difference in Receptor Usage between Severe Acute Respiratory Syndrome (SARS) Coronavirus and SARS-Like Coronavirus of Bat Origin. J Virol. 2008 Feb 15;82(4):1899–907.

22. Ge XY, Li JL, Yang X Lou, Chmura AA, Zhu G, Epstein JH, et al. Isolation and characterization of a bat SARS-like coronavirus that uses the ACE2 receptor. Nature. 2013;503(7477):535–8.

23. Hu B, Zeng LP, Yang X Lou, Ge XY, Zhang W, Li B, et al. Discovery of a rich gene pool of bat SARS-related coronaviruses provides new insights into the origin of SARS coronavirus. PLoS Pathog. 2017; 13(11):e1006698.

24. Lau SKP, Feng Y, Chen H, Luk HKH, Yang W-H, Li KSM, et al. Severe Acute Respiratory Syndrome (SARS) Coronavirus ORF8 Protein Is Acquired from SARS-Related Coronavirus from Greater Horseshoe Bats through Recombination. J Virol. 2015 Oct 15;89(20):10532–47.

25. Ge XY, Wang N, Zhang W, Hu B, Li B, Zhang YZ, et al. Coexistence of multiple coronaviruses in several bat colonies in an abandoned mineshaft. Virol Sin. 2016 Feb 1;31(1):31–40.

